# *De novo* design of modular and tunable allosteric biosensors

**DOI:** 10.1101/2020.07.18.206946

**Authors:** Alfredo Quijano-Rubio, Hsien-Wei Yeh, Jooyoung Park, Hansol Lee, Robert A. Langan, Scott E. Boyken, Marc J. Lajoie, Longxing Cao, Cameron M. Chow, Marcos C. Miranda, Jimin Wi, Hyo Jeong Hong, Lance Stewart, Byung-Ha Oh, David Baker

## Abstract

Naturally occurring allosteric protein switches have been repurposed for developing novel biosensors and reporters for cellular and clinical applications ^1^, but the number of such switches is limited, and engineering them is often challenging as each is different. Here, we show that a very general class of allosteric protein-based biosensors can be created by inverting the flow of information through *de novo* designed protein switches in which binding of a peptide key triggers biological outputs of interest ^2^. Using broadly applicable design principles, we allosterically couple binding of protein analytes of interest to the reconstitution of luciferase activity and a bioluminescent readout through the association of designed lock and key proteins. Because the sensor is based purely on thermodynamic coupling of analyte binding to switch activation, only one target binding domain is required, which simplifies sensor design and allows direct readout in solution. We demonstrate the modularity of this platform by creating biosensors that, with little optimization, sensitively detect the anti-apoptosis protein Bcl-2, the hIgG1 Fc domain, the Her2 receptor, and Botulinum neurotoxin B, as well as biosensors for cardiac Troponin I and an anti-Hepatitis B virus (HBV) antibody that achieve the sub-nanomolar sensitivity necessary to detect clinically relevant concentrations of these molecules. Given the current need for diagnostic tools for tracking COVID-19 ^3^, we use the approach to design sensors of antibodies against SARS-CoV-2 protein epitopes and of the receptor-binding domain (RBD) of the SARS-CoV-2 Spike protein. The latter, which incorporates a *de novo* designed RBD binder, has a limit of detection of 15pM with an up to seventeen fold increase in luminescence upon addition of RBD. The modularity and sensitivity of the platform should enable the rapid construction of sensors for a wide range of analytes and highlights the power of *de novo* protein design to create multi-state protein systems with new and useful functions.

## Main text

Protein-based biosensors play important roles in synthetic biology and clinical applications, but thus far, biosensor design has been mostly limited to reengineering natural proteins ^1^. However, finding analyte-binding domains that undergo sufficient conformational changes is challenging, and even when available, extensive protein engineering efforts are generally required to effectively couple them to a reporter domain ^4, 5^. Hence it is desirable to develop modular biosensor platforms that can be easily repurposed to detect different analytes of interest. Developing modular biosensor platforms from natural analyte-binding proteins, either by creating semisynthetic protein platforms ^6–8^ or based on calmodulin switches ^9, 10^, usually requires random screening to find potential candidates due to limited predictability^11^.

A protein biosensor can be constructed from a system with two nearly isoenergetic states - the equilibrium between which is modulated by the analyte being sensed. Desirable properties in such a sensor are (i) the analyte triggered conformational change should be independent of the details of the analyte (so the same overall system can be used to sense many different compounds) (ii) the system should be tunable so that analytes with different binding energies and relevant concentrations can be detected over a large dynamic range, and (iii) the conformational change should be coupled to a sensitive output. We hypothesized that these attributes could be attained by inverting the information flow in *de novo* designed protein switches in which binding to a target protein of interest is controlled by the presence of a peptide actuator ^2^. As originally described, these switches consist of a constant “cage” region that sequesters a “latch” that binds the target of interest; addition of a peptide “key” displaces the latch from the cage leading to target binding and associated downstream events. However, from a thermodynamic viewpoint, the key and the target are equivalent: the binding of the two to the cage is thermodynamically coupled since the latch has to open, with free energy cost ΔG_open_ (Fig 1b), in order for either to bind. Hence, the free energy associated with binding both target and key is more favorable than the sum of the free energies of binding the two individually (Fig 1c). The difference between key and target is in their variability; the key is constant while the target can be any desired interaction. For an actuator, it is desirable to have a constant input drive a wide range of customizable responses, and hence in our previous work, the input was the (constant) key and the output was binding to a variety of targets associated with protein degradation, nuclear export, etc ^2^. We reasoned that the input to the system could be inverted to create biosensors with a constant readout -- addition of a (variable) target could induce binding of the (constant) key to the (constant) cage, and that this association could be coupled to an enzymatic readout. Such a system would satisfy properties (i) and (ii) above, as a wide range of binding activities can be caged, and since the switch is thermodynamically controlled, it is straightforward to adjust the relative energies of key and target binding to achieve activation at the relevant target concentrations. Because the key and the cage are always the same, the system is modular: the same molecular association can be coupled to the binding of many different targets.

**Fig. 1.**
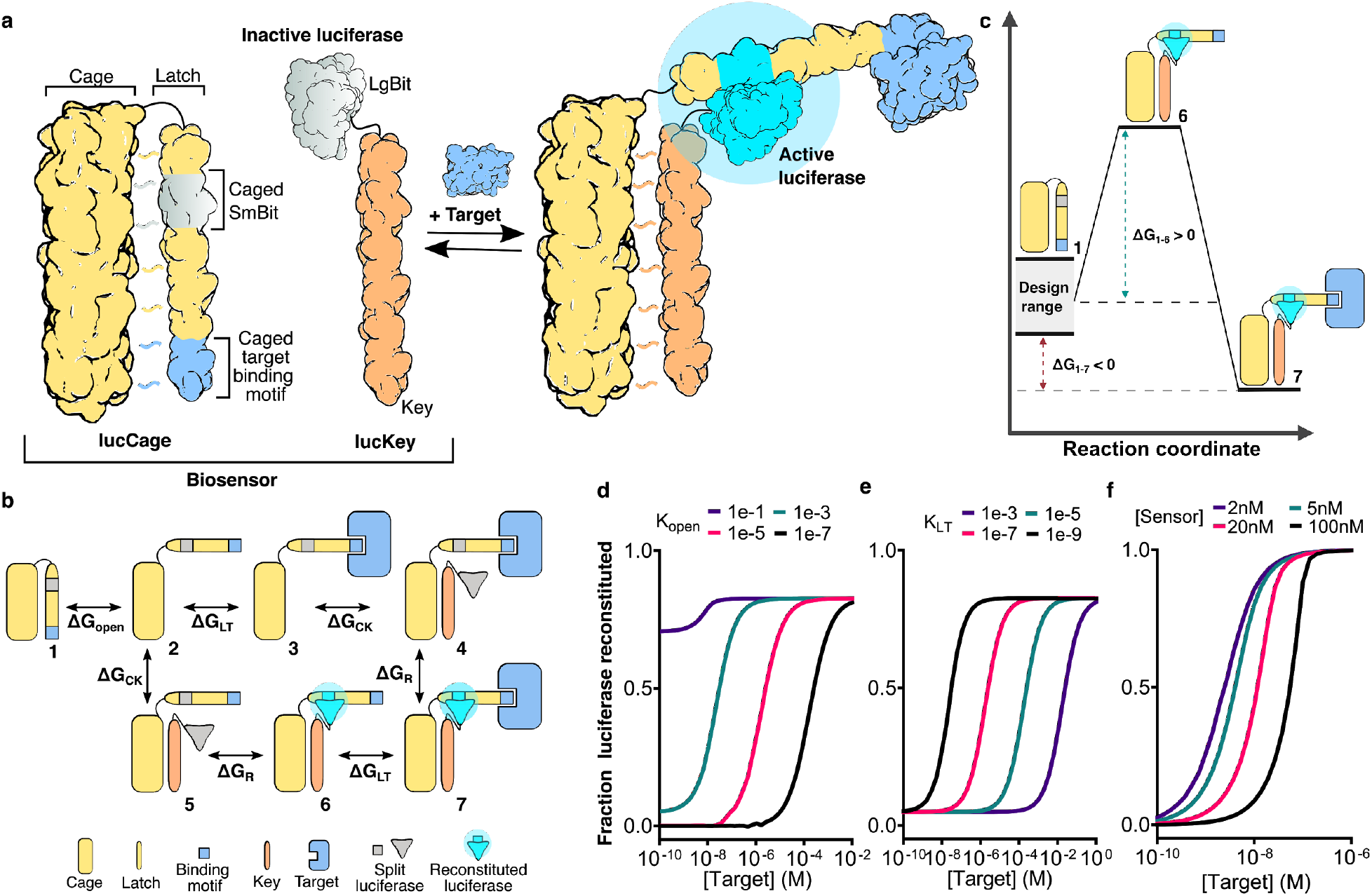
*De novo* design of multi state allosteric biosensors. **a**, Sensor schematic. The biosensor consists of two protein components: lucCage and lucKey, which exist in a closed (Off) and open state (On). The closed form of lucCage (left) can not bind to lucKey, thus, preventing the split luciferase SmBit fragment from interacting with LgBit. The open form (right) can bind both target and key, and allows SmBit to combine with LgBit on lucKey to reconstitute luciferase activity. **b**, Thermodynamics of biosensor activation. The free energy cost ΔG_open_ of the transition from closed cage (species 1) to open cage (species 2) disfavors association of key (species 5) and reconstitution of luciferase activity (species 6) in the absence of target. In the presence of the target, the combined free energies of target binding (2→3; ΔG_LT_), key binding (3→4; ΔG_CK_), and SmBit-LgBit association (4→7; ΔG_R_) overcome the unfavorable ΔG_open_, driving opening of the lucCage and reconstitution of luciferase activity. **c**, Biosensor design strategy based on thermodynamics. For each biosensor, the designable parameters are ΔG_open_ and ΔG_CK_; ΔG_R_ is the same for all targets, and ΔG_LT_ is pre-specified for each target. For sensitive but low background analyte detection, ΔG_open_ and ΔG_CK_ must be designed such that the closed state (species 1) is substantially lower in free energy than the open state (species 6) in the absence of target, but higher in free energy than the open state in the presence of target (species 7). **d-f**, Numerical simulations of the coupled equilibria shown in **b**for different values of (**d**) *K*_open_, (**e**) *K*_LT_, and (**f**) [lucKey]_tot_ and [lucCage]_tot_. *K*_open_, *K*_LT_, *K*_CK_ were set to 1 × 10^−3^, 1 nM, and 10 nM respectively, and the concentration of the sensor components to 10:100 nM (lucCage:lucKey) except where explicitly indicated. **d**, Increasing ΔG_open_ shifts response to higher analyte concentrations. **e**, The sensor limit of detection is approximately 0.1 × *K*_LT_; the driving force for opening the switch becomes too weak below this concentration. **f**, The effective target detection range can be tuned by changing the sensor component concentrations.

To achieve property (iii), we reasoned that bioluminescence could provide a rapid and sensitive readout of analyte driven cage-key association, and explored the use of a reversible split luciferase complementation system ^12^. We developed a system consisting of two protein components: a ‘lucCage’ comprising a cage domain and a latch domain containing the short split luciferase fragment (SmBiT) and an analyte binding motif of choice; and a “lucKey”, which comprises the larger split luciferase fragment (LgBit) and a key peptide (Fig. 1a). lucCage has two states: a closed state in which the cage domain binds the latch and sterically occludes the analyte binding motif from binding its target and SmBiT from combining with LgBit to reconstitute luciferase activity; and an open state in which these binding interactions are not blocked, and lucKey can bind the cage domain. Association of lucKey with lucCage results in the reconstitution of luciferase activity (Fig. 1a, right). The target may be viewed as allosterically regulating luciferase activity, since binding to the sensor is at a site distant from the enzyme active site.

The states of such a system are in thermodynamic equilibrium, with the tunable parameters ΔG_open_ and ΔG_CK_ governing the populations of the possible species, along with the free energy of association of the analyte to the binding domain ΔG_LT_ (Fig. 1b). To achieve high sensitivity, the closed state (species 1) must be substantially lower in free energy than the open state in the absence of target (species 6) to avoid background signal ΔG_1-6>0_), but higher in free energy than the open state in the presence of target (species 7, ΔG_1-7<0_), so that target detection is energetically favorable (Fig. 1c). To guide the optimization of biosensor sensitivity, we simulated the dependence of the sensor system on ΔG_open_ (Fig. 1d), ΔG_LT_ (Fig. 1e), and the concentration of analyte and the sensor components (Fig. 1f) (See Supplementary Methods for details). As expected, the sensitivity of analyte detection is a function of ΔG_LT_, with a lower limit of roughly one-tenth the *K*d for analyte binding (Fig. 1e; below this concentration, the free energy of binding is too small to open the switch). Hence sensing domains with high affinity to their target will yield more sensitive biosensors. The sensitivity of the system can be further tuned above this lower limit by varying the concentration of lucCage and lucKey, resulting in sensing systems responding to different target concentration ranges (Fig. 1f). Tuning the strength of the intramolecular cage-latch interaction (ΔG_open_) affects the equilibrium population of the catalytically active species (species 6 and 7, Fig. 1d), which in turn affects the sensitivity: too tight interaction results in low signal in the presence of target, and too weak an interaction results in high background in the absence of target. Our design strategy aims to find this balance by designing sensors in the closed state (species 1) with a range of ΔG_open_ values: ΔG_open_ can be increased (decreased) by increasing (decreasing) the length of the latch helix and by introducing either favorable hydrophobic interactions or unfavorable steric clashes and buried polar atoms at the cage-latch interface ^2^; we employ both strategies to tune the sensors described below (ΔG_CK_ can also be tuned, but we did not find this necessary for the sensors described here).

To streamline the design of new sensors based on these principles, we developed a Rosetta-based computational method for the incorporation of diverse sensing domains into the LOCKR switches ^2^ called GraftSwitchMover. This method identifies the most suitable position for embedding a target binding peptide within the latch such that the resulting protein is stable in the closed state and the interactions with the target are blocked. This is done by maximizing favorable hydrophobic packing interactions between the peptide and the cage and minimizing the number of unfavorable buried hydrophilic residues. This method takes as input the 3-dimensional model of the switch, the sequence of a peptide that binds the target of interest, and a list of the residues in this peptide that interact with the target (interface residues), and returns a set of designs in which the binding of the peptide to the target is predicted to be blocked by association with the cage (See supplementary methods). The final set of designs covers a range of ΔG_open_values (Fig. 1c), which can be further tuned through introducing destabilizing mutations in the latch: I328S (“1S”) or I328S/L345S (“2S”) ^2^. These designs are then experimentally characterized to find the most sensitive biosensors.

We first set out to test our hypothesis by grafting the SmBiT peptide and the Bim peptide in the closed state of the optimized asymmetric LOCKR switch described in Langan et al, 2020^2^ (Extended Data Fig. 1). SmBiT naturally adopts a β-strand conformation within the luciferase holoenzyme, but we assumed that it will adopt a helical secondary structure in the context of the helical bundle scaffold, consistent with the observation that some peptide sequences can adopt diverse secondary structures in a context-dependent manner ^13^. We sampled different threadings for the two peptide sequences across the latch, built three-dimensional models, selected the lowest energy solutions (3 positions for SmBiT, and 4 positions for the Bim peptide) (Extended Data Fig. 1a) and expressed twelve designs in *E. coli*. We mixed the designs with lucKey in a 1:1 ratio, then added Bcl-2, which binds with nanomolar affinity to Bim, and monitored luciferase activity (Extended Data Fig. 1b). We found that upon the addition of Bcl-2 to a solution containing the new Cage designs, lucKey, and furimazine substrate, there was a rapid increase in luminescence (Extended Data Fig. 1f), suggesting that the inverse LOCKR system can indeed function as a biosensor. Further characterization of the best Bcl-2 sensor candidate, lucCageBim, demonstrated that the analyte detection range could be tuned by varying the concentration of the sensor (lucCage + lucKey) (Extended Data Fig. 1g) as anticipated in our model simulations (Fig. 1f). Experimental characterization of the different designs showed that inserting SmBiT into position 312 of the LOCKR cage (SmBiT312) yielded the highest stability and brightness (Extended Data Fig. 1b), therefore we used this design, henceforward referred to as “lucCage”, as the base scaffold for the biosensors described below.

To explore the versatility of our new biosensor platform, we next investigated the incorporation of a range of binding modalities for analytes of interest within lucCage. First, we set out to explore how to computationally cage target-binding proteins, rather than peptides, in the closed state. We identified the primary interaction surface of the binding protein to its target, extracted the main secondary structure elements involved in it to use them in the computational protocol described above, and selected the best designs from the many threadings generated. Then, we used Rosetta Remodel ^14^ to model the full-length binding domain in the context of the switch and selected designs in which this interface was buried against the cage with minimal steric clashes (See supplementary methods). As a test case, we caged the *de novo* designed protein, HB1.9549.2, which binds to Influenza A H1 hemagglutinin (HA)^15^ into a shortened version of the LOCKR switch (sCage), optimized to improve stability and facilitate crystallization efforts (Fig. 2a). Two of five designs were functional, and bound HA in the presence but not the absence of key (Extended Data Fig. 2b). The crystal structure of the best design, sCageHA_267-1S, determined to 2.0 Å resolution (Table S1), showed that all HA-binding residues except one (F273) interact with the cage domain (blocking binding of the latch to the switch) as intended by design (Fig. 2a, Extended Data Fig. 2a-c). With this structural validation of the design concept in hand, we next sought to develop new sensors using small proteins as sensing domains for the detection of botulinum neurotoxin, the immunoglobulin Fc domain, and the Her2 receptor. To do so, we grafted a *de novo* designed binder for Botulinum neurotoxin B (BoNT/B) ^15^, the C domain of the generic antibody binding protein Protein A ^16^, and a Her2-binding affibody ^17^, into lucCage. After screening a few designs for each target (Extended Data Fig. 3-5), we obtained highly sensitive lucCages (lucCageBot, lucCageProA, and lucCageHer2) that can detect BoNT/B (Fig. 2b, Extended Data Fig. 3), hIgG Fc domain (Fig. 2c, Extended Data Fig. 4), and Her2 receptor (Fig. 2d; Extended Data Fig. 5) respectively, demonstrating the modularity of the platform. The designed sensors responded within minutes upon adding the target, and their sensitivity could be tuned by changing the concentration of lucCage and lucKey (Fig. 2), as predicted by our model simulations (Fig. 1f). Further optimization and characterization of these sensors would enable their use in multiple applications, such as rapid and low-cost detection of highly toxic botulinum neurotoxins in the food industry, which currently relies heavily on live-animal bioassays ^18^, or detection of high serological levels of soluble Her2 (>15 ng/mL) associated with metastatic breast cancer ^19^, levels that could be detected with the current sensitivity of lucCageHer2.

**Fig. 2.**
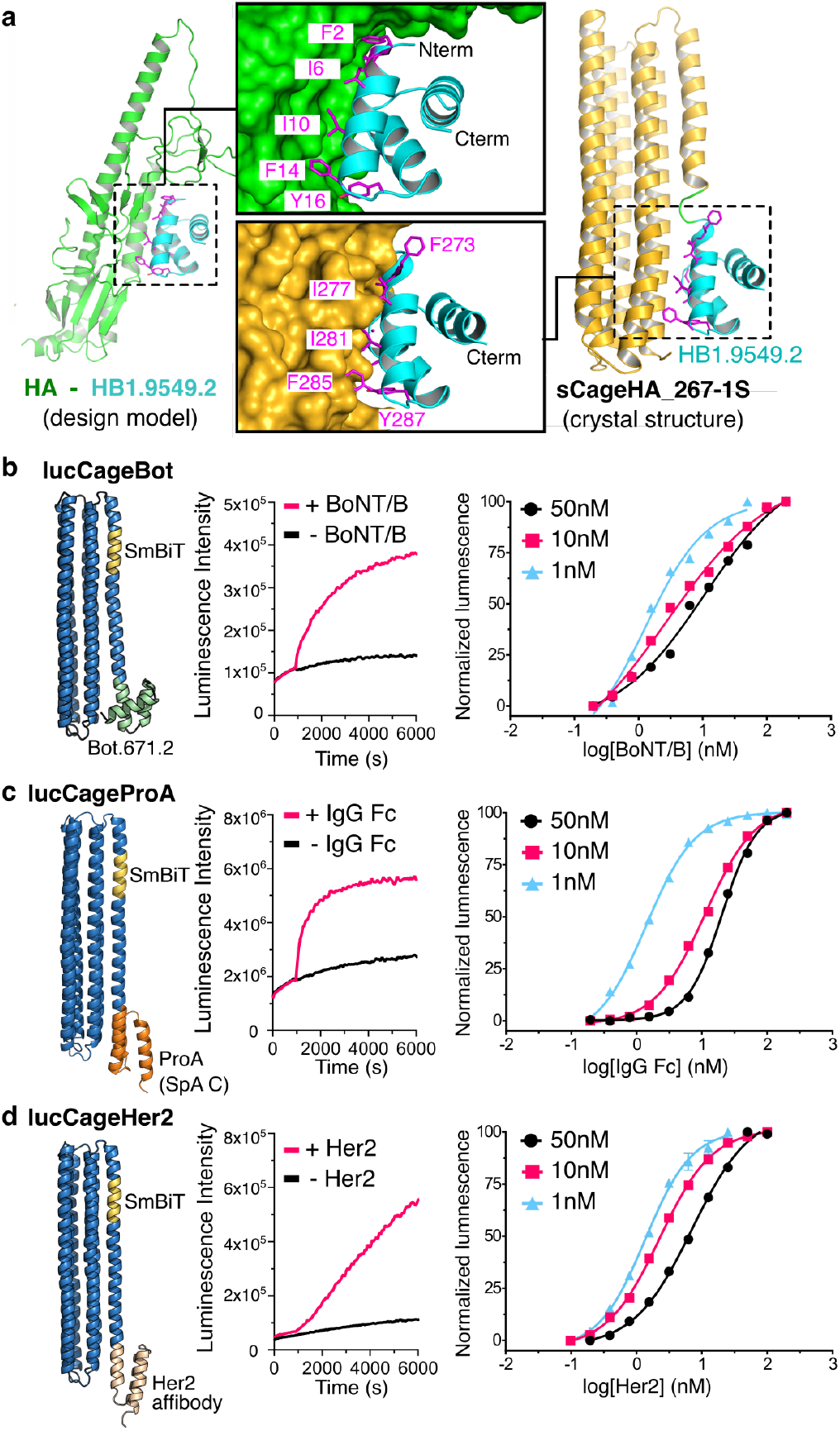
Design and characterization of *de novo* biosensors incorporating small proteins as sensing domains. **a**, General strategy and structural validation for caging small protein domains into LOCKR switches ^2^. Left: design model of the *de novo* binder HB1.9549.2 (cyan) bound to the stem region of influenza hemagglutinin (HA, green ribbon representation) ^15^. Right: crystal structure of sCageHA_267_1S, comprising HB1.9549.2 (cyan) grafted into a shortened and stabilized version of the LOCKR switch (sCage, yellow ribbon representation). Middle: All residues of HB1.9549.2 involved in binding to HA (magenta, top) except for F273 are buried in the closed state of the switch (bottom) to block its interaction. The labels in magenta indicate the same set of amino acids in the two panels (F2 in the top panel corresponds to F273 in the lower panel). **b-d**, Functional characterization of 3 allosteric biosensors: lucCageBot (detection of botulinum neurotoxin B (BoNT/B)), lucCageProA (detection of Fc domain), and lucCageHer2 (detection of Her2 receptor). Left: structural models of the indicated biosensors (ribbon representation) incorporating a *de novo* designed binder for BoNT/B (Bot.671.2) ^15^, the C domain of the generic antibody binding protein Protein A (SpaC) ^16^ and a Her2-binding affibody ^17^ respectively, grafted into lucCage (blue) comprising a caged SmBiT fragment (gold). Middle: kinetic measurement of luminescence intensity upon addition of 50 nM of analyte (BoNT/B, IgG Fc, or Her2) to a mixture of 10 nM of each lucCage and 10 nM of lucKey. Right: detection over a wide range of analyte concentrations by changing the biosensor concentration (50, 5 and 1 nM lucCage and lucKey; cyan, magenta and black lines respectively).

We next designed sensors for additional targets relevant in clinical settings. Since bioluminescent sensors do not require light for excitation, highly sensitive and low background readout is more suited than fluorescence to directly measure analytes in biological media such as blood and serum for point-of-care applications ^20^. We first targeted cardiac troponin I (cTnI), which is the standard early diagnostic biomarker for acute myocardial infarction (AMI). We took advantage of the high-affinity interaction between cTnT, cTnC, and cTnI (Fig. 3a) and designed eleven biosensor candidates by inserting 6 truncated cTnT sequences at different latch positions (Extended Data Fig. 6a). The best candidate, lucCageTrop627, was able to detect cTnI but not at sufficiently low levels for clinical use (Extended Data Fig. 6d). Because the rule-in and rule-out levels of cTnI assay for diagnosis of AMI in patients are in the low pM range ^21^, and because as noted above the limit of detection (LOD) of our sensor platform is about 0.1 x *K*d of the latch-target affinity (*K*_LT_), we further increased the affinity of our sensor to cTnI by fusing cTnC to its terminus (Fig. 3a, Extended Data Fig. 6b,c). The resulting sensor, lucCageTrop, has a single-digit pM LOD suitable for quantification of clinical samples (Fig. 3b, Extended Data Fig. 6e,f).

**Fig. 3.**
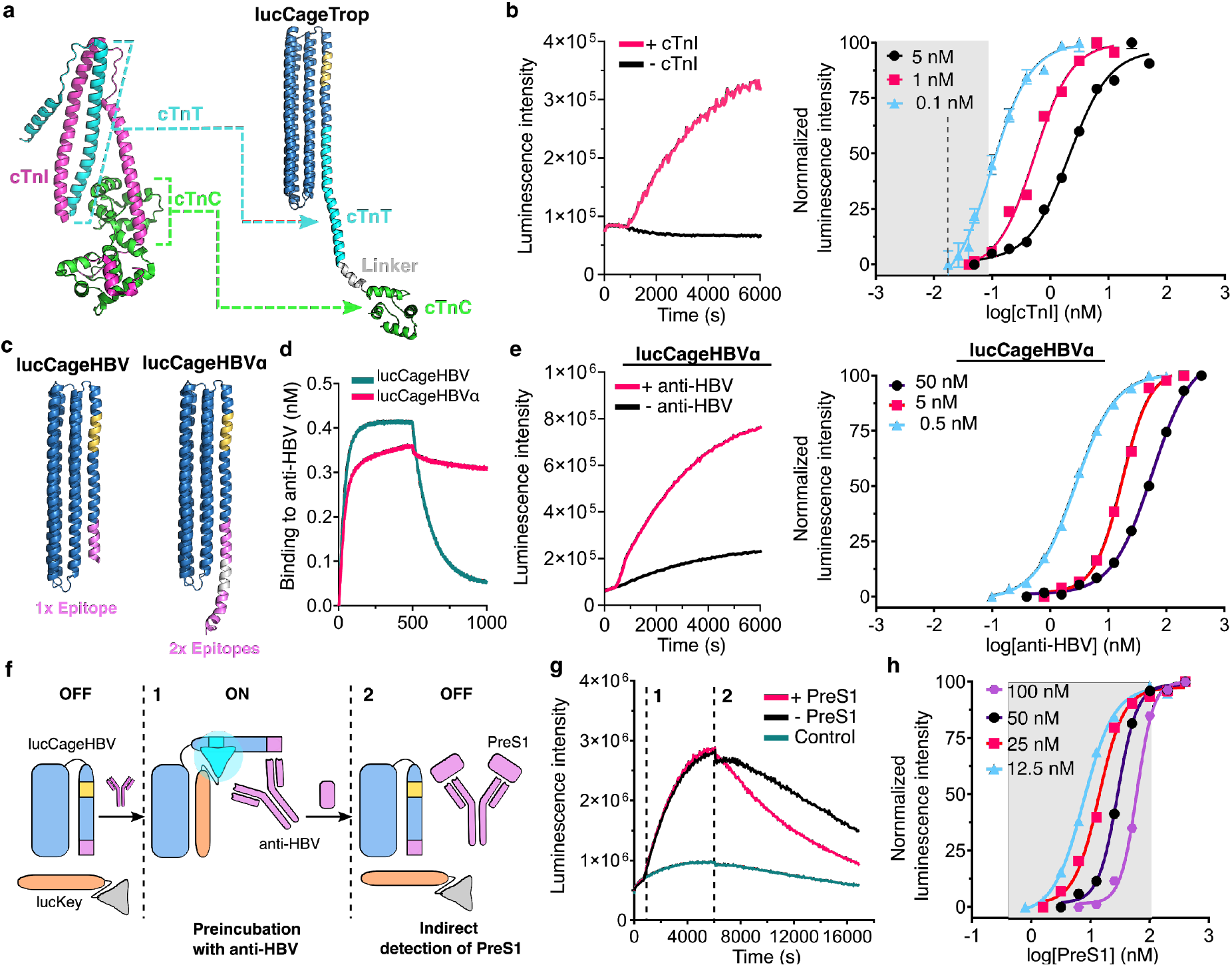
Design and characterization of biosensors for cardiac troponin I and for an anti-HBV antibody. **a**, Design of lucCageTrop, a sensor for cardiac Troponin I. Left: Structure of cardiac troponin (PDB ID: 4Y99); cyan, green, and magenta correspond to cardiac Troponin T (cTnT), cardiac Troponin C (cTnC), and cardiac Troponin I (cTnI), respectively. Right: Design model of lucCageTrop, the cTnI sensor in the closed state containing segments of cTnT (cyan) and cTnC (green). **b**, Left: Kinetics of luminescence increase upon addition of 1 nM cTnI to 0.1 nM lucCageTrop sensor + 0.1 nM of lucKey. Right: A wide analyte (cTnI) detection range can be achieved by changing the concentration of the sensor components (colored lines). The grey area indicates the cTnI concentration range relevant to the diagnosis of acute myocardial infarction (AMI) ^21^; the dotted line indicates clinical AMI cut-off defined by W.H.O. (0.6 ng/mL, 25 pM). **c**, Design models of lucCageHBV and lucCageHBVα, containing SmBit (gold), and one or two tandem antigenic epitopes from the Hepatitis B Virus (HBV) PreS1 protein, respectively (magenta). **d**, lucCageHBVα (two epitope copies) has higher affinity for the anti-HBV antibody HzKR127-3.2 (*K*d= 0.68 nM) than lucCageHBV (one epitope copy) (Kd= 20 nM) as demonstrated by biolayer interferometry. **e**, Left: Kinetics of bioluminescence signal increase upon addition of 10n anti-HBV antibody to 1nM lucCageHBVα + 1nM lucKey. Right: By varying the concentrations of the sensor components, sensitive anti-HBV antibody detection can be achieved over a wide concentration range. **f**, Schematic of the detection mechanism for HBV protein PreS1 using lucCageHBV. **g**, Kinetics of bioluminescence following addition of the anti-HBV antibody (step 1) and subsequently PreS1 (step 2). The bioluminescence decreases upon PreS1 addition as PreS1 competes with the sensor for the antibody. **h**, Sensitive detection of PreS1 can be achieved over the relevant post-HBV infection concentration levels (grey area). The sensor is pre-mixed with the anti-HBV antibody; the PreS1 detection range can be tuned by varying the concentration of antibody (indicated by colored labels).

Detection of specific antibodies is important for monitoring the spread of a pathogen in a population (antibodies remain long after the pathogen has been eliminated) ^22^, the success of vaccination ^23^, and levels of therapeutic antibodies ^24^. To adapt our system to be used in such antibody serological analyses, we sought to incorporate linear epitopes recognized by the antibodies of interest into lucCage, so that binding of an antibody would open the switch allowing lucKey binding and reconstitution of luciferase activity. We first developed a sensor for anti-Hepatitis B virus (HBV) antibodies based on the crystal structure of the neutralizing antibody (HzKR127) bound to a peptide from the PreS1 domain of the viral surface protein L ^25^. The best of 8 designs tested, lucCageHBV (HBV344), had a ~150% increase in luciferase activity upon addition of HzKR127-3.2, an improved version of HzKR127 ^26^ (Extended Data Fig. 7a,b). To further improve the dynamic range and LOD of lucCageHBV (~2 nM, Extended Data Fig. 7c-e), we increased the latch-target affinity (*K*_LT_) by introducing an additional copy of the peptide at the end of the latch to take advantage of the antibody bivalent interaction with its epitope (Fig. 3c,d). The resulting design, named lucCageHBVα, had a LOD of 260 pM and a dynamic range of 225% (Fig. 3e; Extended Data Fig. 8a-c), with a luminescence intensity easily detectable with a camera (Extended Data Fig. 8d). Hence the platform to detect specific antibodies with a LOD in the range for monitoring therapeutic antibodies ^27^. We next demonstrated the use of the lucCageHBV sensor to detect hepatitis B surface antigen (HBsAg). Since our sensors are under thermodynamic control, we hypothesized that the pre-assembly of sensor-antibody complex would re-equilibrate in the presence of the target HBsAg protein, PreS1, with antibody redistributing to bind free PreS1 instead of the epitope on lucCageHBV (Fig. 3f). Indeed, the luminescence of lucCageHBV plus HzKR127-3.2 mixture decreased shortly upon addition of the PreS1 domain (Fig. 3g); the sensitivity of this readout enabled quantification of PreS1 concentration in a clinically relevant range^28^ (Fig. 3h, Extended Data Fig. 7f). HBsAg seroclearance is one of the major biomarkers to monitor therapeutic progress following hepatitis diagnosis ^29^ and vaccination efficacy, but current commercial HBsAg assays are unable to differentiate between the three HBsAg protein subtypes. While we have not yet generated sensors for all three subtypes, our PreS1 sensor (detecting HBsAg L antigen) shows that the system can achieve subtype-specific recognition.

The COVID-19 pandemic has showcased the urgent need for developing new diagnostic tools for tracking active infections by detecting the SARS-CoV-2 virus itself, and for detection of antiviral antibodies to evaluate the extent of the spread of the virus in the population and to identify individuals at lower risk of future infection ^30^. To design sensors for anti-SARS-CoV-2 antibodies, we first identified from the literature highly immunogenic linear epitopes in the SARS-CoV ^31, 32^ and SARS-CoV-2 proteomes ^33, 34^ that are not present in “common” strains of *coronaviridae* (i.e., HCoV-OC43, HCoV-HKU1, HCoV-229E, HCoV-NL63; we did not exclude reactivity against SARS-CoV or MERS as they are much less broadly distributed). Among these, we focused on two epitopes in the Membrane and Nucleocapsid proteins found to be recognized by SARS and COVID-19 patient sera for which cross-reactive animal-derived antibodies are commercially available (see Fig. 4 legend and Materials and methods for epitope and antibody description). We designed sensors for each epitope (Extended Data Fig. 9a,b) and identified designs that specifically responded to the presence of pure anti-M and anti-N protein antibodies (Fig. 4b,c). These sensors were fast (2-5 minutes to reach full signal) and had a ~50-70% dynamic range in response to low nanomolar amounts of antibodies, consistent with our previous findings (Fig. 4b,c, Extended Data Fig. 9c,d).

**Fig. 4.**
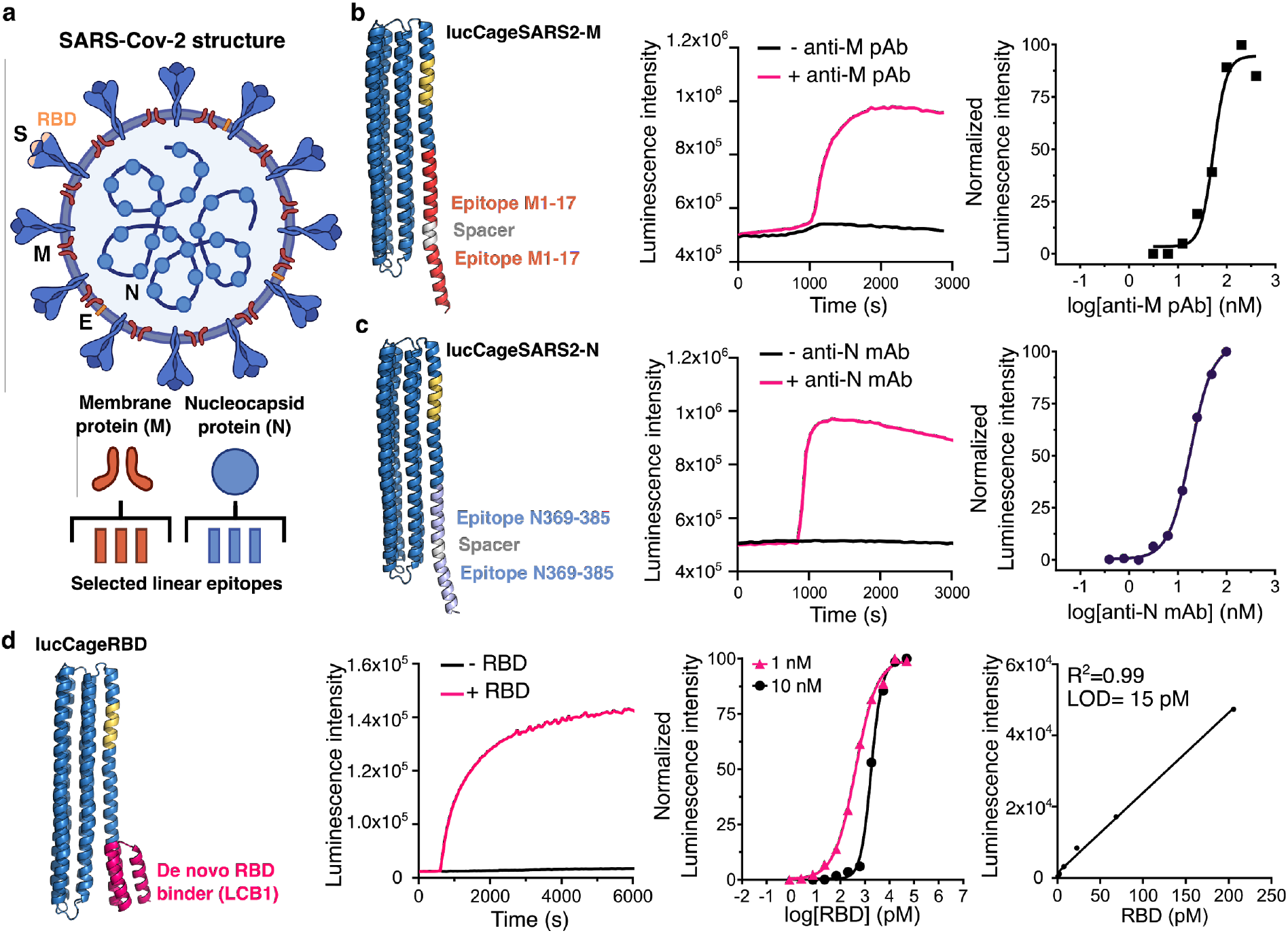
Design of biosensors for detection of anti-SARS-CoV-2 antibodies and SARS-CoV-2 RBD. **a**, SARS-CoV-2 viral structure representation showing the major structural proteins: Envelope protein (E), membrane protein (M), nucleocapsid protein (N), and the Spike protein (S) containing the receptor-binding domain (RBD). Linear epitopes for the M and N proteins were selected based on published immunogenicity data. **b**, Left panel: structural model of lucCageSARS2-M. Two copies of the SARS-CoV-2 Membrane protein a.a. 1-17 epitope (red) are grafted into lucCage connected with a flexible spacer. Middle panel: kinetics of luminescent activation of lucCageSARS2-M (50 nM) + lucKey (50nM) upon addition of anti-SARS-CoV-1 Membrane protein rabbit polyclonal antibodies at 100 nM (ProSci, 3527). These antibodies, originally raised against a peptide corresponding to 13 amino acids near the amino-terminus of SARS-CoV Matrix protein, cross-react with residues 1-17 of the SARS-CoV-2 Membrane protein. Right panel: response of lucCageSARS2-M (5 nM) + lucKey (5nM) to varying concentrations of target anti-M pAb. **c**, Left panel: structural model of lucCageSARS2-N. Two copies of the SARS-CoV-2 Nucleocapsid protein 369-382 epitope (lightblue) are grafted into lucCage connected with a flexible spacer. Middle panel: kinetics of luminescent activation of lucCageSARS2-N (50 nM) + lucKey (50nM) upon addition of 100 nM anti-SARS-CoV-1-N mouse monoclonal antibody (clone 18F629.1). This antibody originally raised against residues 354-385 of the SARS-CoV-1 Nucleocapsid protein cross-reacts with residues 369-382 of the SARS-CoV-2 Nucleocapsid protein. Right panel: response of lucCageSARS2-N (50 nM) + lucKey (50nM) to varying concentration of target (anti-N mAb). **d**, Functional characterization of lucCageRBD, a SARS-CoV-2 RBD sensor. Left panel: structural model of lucCageRBD showing the LCB1 binder (magenta) grafted into lucCage (blue) comprising a caged SmBiT fragment (gold). Second panel: kinetic measurement of luminescence intensity upon addition of 16.7 nM of RBD to a mixture of 1 nM of lucCageRBD and 1 nM of lucKey. Third panel: detection over a wide range of analyte concentrations by changing the biosensor concentration (10 and 1 nM lucCage and lucKey; black and magenta lines respectively). Right panel: Limit of detection (LOD) determination of lucCageRBD and lucKey at 1 nM each for detection of RBD in solution. LOD was determined to be 15 pM.

To create sensors capable of detecting SARS-CoV-2 viral particles directly, we integrated into the LucCage format a designed picomolar affinity binder to the receptor-binding domain (RBD) of the SARS-CoV-2 Spike protein named LCB1 (Fig. 4d). Of 13 candidates tested, the best, which we refer to as lucCageRBD, had minimal background, an outstanding dynamic range (1700%) easily detectable with a camera and low LOD (15 pM) (Fig. 4d, Extended Data Fig. 10). The superior dynamic range and sensitivity of this sensor are consequences of the high affinity of LCB1 to RBD (K_LT_), consistent with our thermodynamic model, highlighting the synergy of the LucCage sensor platform and *de novo* binder design.

Because of the modularity and engineerability of the LucCage system, it took only three weeks to design the SARS-CoV-2 antibody and RBD sensors, obtain synthetic genes, express and purify the proteins, and evaluate sensor performance. There are challenges still to overcome to deploy these in the current pandemic. For serological analysis, generation of an expanded set of sensors spanning the epitopes recognized by infected patient sera will be necessary, and for direct virus detection, characterization and optimization of the sensitivity for live virus particles (with ~100 RBDs per particle, we do not expect the LOD for virus particles to be less than 100 fM; like other protein based sensors lucCageRBD cannot achieve the near single virion detection sensitivity of nucleic acid amplification based methods).

To test the specificity of the biosensors developed in this work (excluding the indirect detection of PreS1 by lucCageHBV), we measured the activation kinetics of each in response to all the targets (Bcl-2, botulinum neurotoxin B, IgG Fc, Her2, cardiac Troponin I, the monoclonal anti-HBV antibody (HzKR127-3.2), the anti-SARS-CoV-1-M polyclonal antibody (clone 3527), the anti-SARS-CoV-1-N monoclonal antibody (clone 18F629.1), and PreS1). As shown in Fig. 5, each sensor responded rapidly and sensitively to its cognate target, but not to any of the others. A summary of each lucCage sensor characteristics and sensing domains used can be found in Table S2 and Table S3, respectively.

**Fig. 5.**
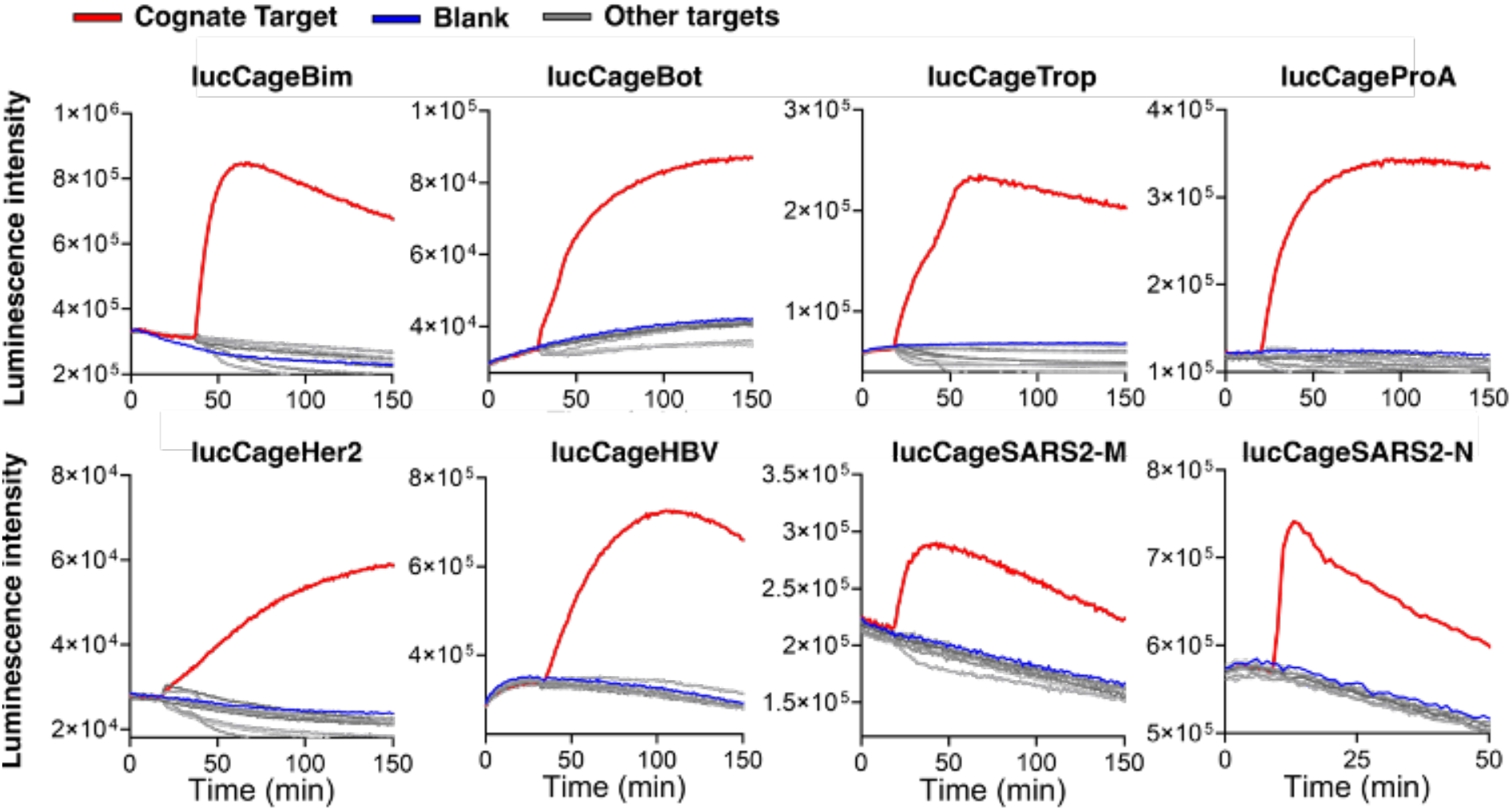
Biosensor specificity. Each sensor at 10 nM (LucCageSARS2-N at 50nM) was incubated with 50 nM of its cognate target (red lines), the targets for the other biosensors (grey) or buffer (blue). Targets are Bcl-2, botulinum neurotoxin B, IgG Fc, Her2, cardiac Troponin I, the monoclonal anti-HBV antibody (HzKR127-3.2), the anti-SARS-CoV-1-M polyclonal antibody (3527), the anti-SARS-CoV-1-N monoclonal antibody (clone 18F629.1), SARS-CoV-2 RBD and PreS1. Strong responses were observed only for the cognate targets.

It is instructive to put our sensors in the context of the multiple protein-based biosensor platforms that have been developed over the years to detect small molecules ^20, 35^, antibodies ^7, 24^, and other protein ligands ^6^ with considerable success (see Supplementary discussion section, Extended Data Fig. 11, Table S4). Most of these platforms depend on the specific geometry of a target-sensor interaction to trigger a conformational change in the reporter component and hence are specialized for a subset of detection challenges. Because of this target dependence, considerable optimization can be required to achieve high sensitivity detection of a new target. Our sensor platform is based on the thermodynamic coupling between defined closed and open states of the system, thus, its sensitivity depends on the free energy change upon the sensing domain binding to the target but not the specific geometry of the binding interaction (the semi-synthetic small molecule sensors ^20, 35^ also have this property). This enables the incorporation of various binding modalities, including small peptides, globular mini proteins, antibody epitopes and *de novo* designed binders, to generate sensitive sensors for a wide range of protein targets with little or no optimization. For point of care (POC) applications, our system, like other protein biosensor platforms ^36^, has the advantages of being homogeneous, no-wash, all-in-solution, a nearly instantaneous readout, and its quantification of luminescence could be performed by means of inexpensive and accessible devices such as a cell phone camera. In hospital settings, the ability to predictably make a wide range of sensors under the same principle could enable quick readout of large numbers of different compounds using an array of hundreds of different sensors on, for example, a 384-well plate. However, to achieve optimal performance comparable to the best of the previously described approaches, e.g., the LUMAB antibody detection system^36^, the sensors described here will likely require further engineering by fine-tuning the thermodynamic parameters outlined in Fig. 1. Still to be worked out are the most effective ways to accurately quantify analyte presence in complex biological fluids where the luminescence readout could be disturbed, such as blood, serum, or saliva. Future modifications of the sensor architecture, such as detecting bioluminescence resonance energy transfer (BRET) rather than direct luminescence ^37^ could increase the accuracy of quantification at a cost of some sensitivity loss.

Up until recently, the focus of *de novo* protein design was on the design of proteins with new structures corresponding to single deep free energy minima; our results highlight the progress in the field which now enables more complex multistate systems to be readily generated. Our sensors, like other *de novo* designed proteins, are expressed at high levels in cells and are very stable, which should considerably facilitate the further manufacturing process. The general “molecular device” architecture of our platform synergizes particularly well with complementary advances in the *de novo* design of high-affinity miniprotein binders ^15^, which can be designed with three dimensional structures readily compatible with the lucCage platform (designed binders has also been incorporated into the LUMAB antibody detection platform ^36^). LucCageRBD highlights the potential of this fully *de novo* approach, with a 1700% dynamic range and 15 pM LOD from a sensor coming straight out of the computer, without any experimental optimization. As the power of computational design continues to improve, it should become possible to detect an ever wider range of targets with greater sensitivity using LucCage sensors. Beyond biosensors, our results highlight the potential of *de novo* protein design to create more general solutions for current day challenges than can be achieved by repurposing native proteins that have evolved to solve completely different challenges.

## Funding

We acknowledge funding from HHMI (D.B.), the LG Yonam Foundation (B.-H.O.), the National Research Foundation (B.-H.O., NRF-2018R1A2B3004764), the BK21 PLUS project of Korea (H.L.), the United World Antiviral Research Network (UWARN one of the Centers Researching Emerging Infectious Diseases “CREIDs”, NIAID 1 U01 AI151698-01, D.B. and L.S.), The Audacious Project at the Institute for Protein Design (D.B., C.M.C., and M.C.M.), Eric and Wendy Schmidt by recommendation of the Schmidt Futures (A.Q-.R. and H-.W.Y.), the Washington Research Foundation (J.P. and M.J.L.), the Nordstrom Barrier Institute for Protein Design Directors Fund (R.A.L.), and The Open Philanthropy Project Improving Protein Design Fund (D.B. and S.E.B.), the gift support from Gree Real Estate (A.Q.-R.), and “la Caixa” Foundation (A.Q.-R., ID 100010434 under grant LCF/BQ/AN15/10380003).

We thank Dr. Wesley C. Van Voorhis for advice and support with the anti-SARS-CoV-2 antibody sensors, Nuttada Panpradist and Dr. Barry Lutz for providing simulated nasal matrix, Stephanie Berger for sharing the Bcl2 protein target, and Alex Kang for setting up screening crystal trays. The X-ray data were collected on the Beamline 5C at the Pohang Accelerator Laboratory, Korea. All structural images for figures were generated using PyMOL 2.0. Some figures in this article were created with Biorender.com.

## Data availability

The coordinates of the structure of sCageHA_267-1S will be deposited in the Protein Data Bank with the conditions of immediate release upon publication. The original data that support the findings are available from the corresponding authors upon reasonable request. Plasmids encoding the biosensor proteins described in this article are available from the corresponding authors upon reasonable request.

## Code availability

Relevant code used in this manuscript will be deposited in a public repository upon publication.

## Author contributions

D.B. directed the work. D.B., A.Q.-R., B.-H.O., H.-W.Y., J.P. and L.S. designed and further conceptualized the research. A.Q.-R., B.-H.O., H.-W.Y. and J.P. performed the computational design of the sensors and their experimental validation. B.H.O. directed, and H.L. performed the crystallographic work. R.A.L and H.-W.Y. wrote the thermodynamic model and performed the simulations. R.A.L wrote GraftSwitchMover. S.E.B. and M.J.L. designed the parental cage and key protein scaffolds. L.C. designed the RBD binder. C.M.C., M.C.M., J.W. and H.J.H. performed production and purification of proteins. D.B, B.-H.O., A.Q.-R. and H.-W.Y. wrote the original draft. All authors reviewed and accepted the manuscript.

## Competing interests

D.B., A.Q.-R., H.-W. Y., J.P., B.-H. O., L.C., and L.S. are co-inventors in a provisional patent application covering the biosensors described in this article.

**Correspondence and requests for materials** should be addressed to D.B. or B.-H.O.

## Methods

### Design of the sensor system: lucCage and lucKey

SmBit (VTGYRLFEEIL)^12^ was grafted into the latch of the asymmetric LOCKR switch described in Langan et al, 2019 ^2^ using GraftSwitchMover, a RosettaScripts-based protein design algorithm (See Supplementary Methods for details). The grafting sampling range was assigned between residues 300-330. The resulting designs were energy-minimized, visually inspected and selected for subsequent gene synthesis, protein production and biochemical analyses. The best SmBit position on the latch was experimentally determined to be an insertion at residue 312, as described in Extended Data Fig. 1. lucKey was assembled by genetically fusing the LgBit of NanoLuc ^12^ to the key peptide described in Langan et al, 2019. (See Table S5 for the full sequence list)

### Computational grafting of sensing domains into lucCage

#### Peptides and epitopes

The amino acid sequence for each sensing domain was grafted using Rosetta GraftSwitchMover into all α-helical registers between residues 325-360 of lucCage (See Supplementary Methods for details). The resulting lucCages were energy-minimized, visually inspected and typically less than ten designs were selected for subsequent protein production and biochemical characterization.

#### Protein domains

First, the main secondary structure elements forming the interaction surface of the binding protein were identified, their amino acid sequence was extracted and grafted into lucCage using theGraftSwitchMover as described above. Then, we used Rosetta Remodel ^14^ to model the full-length binding domain in the context of the switch in which this interface was buried against the cage (See Supplementary Methods for details). The designs were energy-minimized and visually inspected for selection. Typically, less than ten designs were selected for biochemical characterization.

### Synthetic gene construction

The designed protein sequences were codon optimized for *E. coli* expression (IDT codon optimization tool) and ordered as synthetic genes in pET21b+ or pET29b+ *E. coli* expression vectors (IDT). The synthetic gene was inserted at the NdeI and XhoI sites of each vector, including an N-terminal hexahistidine tag followed by a TEV protease cleavage site and a stop codon was added at the C terminus.

### General procedures for bacterial protein production and purification

The *E. coli* LEMO21(DE3) strain (NEB) was transformed with a pET21b+ or pET29b+ plasmid encoding the synthesized gene of interest. Cells were grown for 24 hours in LB media supplemented with carbenicillin or kanamycin. Cells were inoculated at a 1:50 mL ratio in the Studier TBM-5052 autoinduction media supplemented with carbenicillin or kanamycin, grown at 37 °C for 2-4 hours, and then grown at 18 °C for an additional 18 h. Cells were harvested by centrifugation at 4000*g* at 4 °C for 15 min and resuspended in 30 ml lysis buffer (20 mM Tris-HCl pH 8.0, 300 mM NaCl, 30 mM imidazole, 1 mM PMSF, 0.02 mg/mL DNAse). Cell resuspensions were lysed by sonication for 2.5 minutes (5 second cycles). Lysates were clarified by centrifugation at 24,000*g*at 4 °C for 20 min and passed through 2 ml of Ni-NTA nickel resin (Qiagen, 30250) pre-equilibrated with wash buffer, (20 mM Tris-HCl pH 8.0, 300 mM NaCl, 30 mM imidazole). The resin was washed twice with 10 column volumes (CV) of wash buffer, and then eluted with 3 CV of elution buffer (20 mM Tris-HCl pH 8.0, 300 mM NaCl, 300 mM imidazole). The eluted proteins were concentrated using Ultra-15 Centrifugal Filter Units (Amicon) and further purified by using a Superdex^TM^ 75 Increase 10/300 GL (GE Healthcare) size exclusion column in Tris Buffered Saline (TBS; 25 mM Tris-HCl pH 8.0, 150 mM NaCl). Fractions containing monomeric protein were pooled, concentrated, and snap-frozen in liquid nitrogen and stored at −80 °C.

### *In vitro* bioluminescence characterization

A Synergy Neo2 Microplate Reader (BioTek) was used for all in vitro bioluminescence measurements. Assays were performed in 1:1=HBS-EP:Nano-Glo assay buffer for anti-HBV and RBD sensors while 1:1=DPBS:Nano-Glo assay buffer was used for other sensors. 10X lucCage, 10X lucKey, and 10X target proteins of desired concentrations were first prepared from stock solutions. For each well of a white opaque 96-well plate, 10 μL of 10X lucCage, 10 μL of 10X lucKey, and 20 μL of buffer were mixed to reach the indicated concentration and ratio. The plate was centrifuged at 1000 × g for 1 min and incubated at RT for additional 10 min. Then, 50 μL of 50X diluted furimazine (Nano-Glo luciferase assay reagent, Promega) was added to each well. Bioluminescence measurements in the absence of target were taken every 1 min post-injection (0.1 s integration and 10 s shaking during intervals). After ~15 min, 10 μL of serially diluted 10X target protein plus a blank was injected and bioluminescence kinetic acquisition continued for a total of 2 h. To derive EC_50_ values from the bioluminescence-to-analyte plot, the top three peak bioluminescence intensities at individual analyte concentrations were averaged, subtracted from blank, and used to fit the sigmoidal 4PL curve. To calculate the LOD, the linear region of bioluminescence responses of sensors to its analyte was extracted and a linear regression curve was obtained. It was used to derive the standard deviation of the response (SD) and the slope of the calibration curve (S). The LOD was determined as 3×(SD/S). The experimental measurements were taken in triplicate and the mean values are shown where applicable. The results were successfully replicated using different batches of pure proteins on different days.

### Biolayer interferometry (BLI)

Protein-protein interactions were measured by using an Octet^®^ RED96 System (ForteBio) using streptavidin-coated biosensors (ForteBio). Each well contained 200 μL of solution, and the assay buffer was HBS-EP+ Buffer (10 mM HEPES pH 7.4, 150 mM NaCl, 3 mM EDTA, 0.05% v/v Surfactant P20, 0.5% non-fat dry milk). The biosensor tips were loaded with analyte peptide/protein at 20 μg/mL for 300 s (threshold of 0.5 nm response), incubated in HBS-EP+ Buffer for 60 s to acquire the baseline measurement, dipped into the solution containing Cage and/or Key for 600 s (association step) and dipped into the HBS-EP+ Buffer for 600 s (dissociation steps). The binding data were analyzed with the ForteBio Data Analysis Software version 9.0.0.10.

### Design and characterization of lucCageBim

The Bim peptide sequence (EIWIAQELRRIGDEFNAYYAAA) was threaded into the lucCage scaffold as described in the “Design of sensing domains into lucCage” section. The selected designs were expressed in *E. coli*, purified and characterized for luminescence activation. The bioluminescence detection signal was measured for each design lucCage at 20 nM mixed with lucKey at 20 nM, in the presence or absence of target Bcl-2 protein at 200nM. Bcl-2 was expressed as described somewhere else ^40^.

### Design and characterization of lucCageHer2, lucCageProA, lucCageBot and lucCageRBD

The main binding motifs of the Bot.0671.2 de novo binder, *S. aureus* Protein A domain C (SpaC), the Her2 affibody and the *de novo* RBD binder LCB1 were threaded into lucCage as described in the “Design of sensing domains into lucCage” section (See Table S3 for sequences of sensing domains). The selected designs were expressed in *E. coli*, purified and characterized for luminescence activation. The bioluminescence detection signal was measured for each design lucCage at 20 nM mixed with lucKey at 20 nM, in the presence or absence of 200nM target protein. The target proteins used were: Botulinum Neurotoxin B HcB expressed as previously described ^41^, human IgG1 Fc-HisTag (AcroBiosystems, Cat. No. IG1-H5225) and human Her2-HisTag (AcroBiosystems, Cat. No. HE2-H5225).

### Design and characterization of lucCageTrop

The cardiac Troponin T (cTnT) binding motif (EDQLREKAKELWQTIYNLEAEKFDLQEKFKQQKYEINVLRNRINDNQ) was split into fragments of different length (see Extended Data Fig. 6) and threaded into the lucCage scaffold as described in the “Design of sensing domains into lucCage” section. The selected designs were expressed in *E. coli*, purified and characterized for luminescence activation. The bioluminescence detection signal was measured for each design lucCage at 20 nM mixed with lucKey at 20 nM in the presence or absence of 100 nM cardiac Troponin I (Genscript, Cat. No. Z03320-50). Subsequently, lucCageTrop, an improved version by fusion to cardiac Troponin C (cTnC), was created by genetically fusing the following sequence to the C terminus of lucCageTrop627 (KVSKTKDDSKGKSEEELSDLFRMFDKNADGYIDLEELKIMLQATGETITEDDIEELMKDGDKNNDGRIDYDEFLEFMKGVE).

### Design and characterization of lucCageHBV and lucCageHBVα

The binding motif (GANSNNPDWDFN) of the PreS1 domain was threaded into the lucCage scaffold at every position after residues 336 using the Rosetta GraftSwitchMover. Following the Rosetta FastRelax protocol, eight designs were selected for protein production. Bioluminescence was measured with the designed lucCages (20 nM) and lucKey (20 nM) in the presence or absence of the anti-HVB antibody HzKR127-3.2 (100 nM) to select lucCageHBV. Subsequently, lucCageHBVα was constructed by genetically fusing a sequence containing a second antigenic motif (GGSGGGSSGF<u>GANSNNPDWDFN</u>PN) to lucCageHBV.

### Design and characterization of lucCageSARS2-M and lucCageSARS2-N

Antigenic epitopes of the SARS-CoV-2 membrane protein (a.a. 1-31, 1-17 and 8-24) and the nucleocapsid protein (a.a. 368-388 and 369-382) were computationally grafted into lucCage as described in the “Design of sensing domains into lucCage” section. The selected designs were expressed in *E. coli*, purified and characterized for luminescence activation. All designs at 50nM were mixed with 50nM lucKey and experimentally screened for an increase in luminescence in the presence of rabbit anti-SARS-CoV Membrane polyclonal antibodies (ProSci, Cat. No.: 3527) at 100nM or mouse anti-SARS-CoV Nucleocapsid monoclonal antibody (clone 18F629.1, NovusBio Cat. No. NBP2-24745) at 100 nM.

### Design and characterization of sCageHA variants

HB1.9549.2 was embedded into the parental six-helix bundle for sCage design at different positions along the latch helix of the scaffold. To promote more favorable intramolecular interactions, three consecutive residues on the latch were intentionally substituted with glycine to allow for conformational freedom. The five designs were produced in *E. coli*. Biolayer interferometry analysis was performed with purified Cages (1 μM) and biotinylated Influenza A H1 hemagglutinin (HA)^15^ loaded onto streptavidin-coated biosensor tips (ForteBio) in the presence or absence of the key (2 μM) using an Octet instrument (ForteBio).

### Production and purification of HzKR127-3.2

The synthetic V_H_ and V_L_ DNA fragments were subcloned into the pdCMV-dhfrC-cA10A3 plasmid containing the human Cγ1 and C*κ* DNA sequences. The vector was introduced into HEK 293T cells using Lipofectamine (Invitrogen), and the cells were grown in FreeStyle 293 (GIBCO) in 5% CO_2_ in a 37 °C humidified incubator. The culture supernatant was loaded onto a protein A-sepharose column (Millipore), and the bound antibody was eluted by the addition of 0.2 M glycine–HCl (pH 2.7), followed by immediate neutralization with 1 M Tris–HCl (pH 8.0). The solution was dialyzed against 10 mM HEPES-NaOH (pH 7.4), and the purity of the protein was analyzed by SDS-PAGE.

### Production and purification of the PreS1 domain

The DNA fragment encoding the PreS1 domain (residues 1-56) was cloned into the pGEX-2T (GE Healthcare) plasmid, and the protein was produced in the *E. coli* BL21(DE3) strain (NEB) at 18 ^o^C as a fusion protein with glutathion-S-transferase (GST) at the N-terminus. The cell lysates were prepared in a buffer solution (25 mM Tris-HCl pH 8.0, 300 mM NaCl), and clarified supernatant was loaded onto GSTBind™ Resin (Novagen). The GST-PreS1 domain was eluted with the same buffer containing additional 10 mM reduced glutathione, further purified using a Superdex^TM^ 75 Increase 10/300 GL (GE Healthcare) size exclusion column, and concentrated to 34 μM.

### Production of SCageHA_267-1S and its variants

sCageHA_267-1S and sCageHA_267-1S(E99Y/T144Y) were expressed at 18 °C in the *E. coli* LEMO21(DE3) strain (NEB) as a fusion protein containing a (His)10-tagged cysteine protease domain (CPD) derived from *Vibrio cholerae* ^42^ at the C-terminus. The protein was purified using HisPur^TM^ nickel resin (Thermo), a HiTrap Q anion exchange column (GE Healthcare) and a HiLoad 26/60 Superdex 75 gel filtration column (GE Healthcare). For Selenomethionine (SelMet)-labeling, an I30M mutation was introduced additionally to generate a sCageHA_267-1S(E99Y/T144Y/I30M) variant. This protein was expressed in the *E. coli* B834 (DE3) RIL strain (Novagen) in the minimal media containing SeMet, and purified according to the same procedure for purifying the other variants.

### Crystallization and structure determination of sCageHA_267-1S

Two point mutations (Glu99Tyr and Thr144Tyr) were introduced in an attempt to induce favorable crystal packing interactions. Good-quality single crystals of sCageHA_267-1S(E99Y/T144Y/I30M) were obtained in a hanging-drop vapor-diffusion setting by micro-seeding in a solution containing 11% (v/v) ethanol, 0.25 M NaCl, 0.1 M TrisHCl (pH 8.5). The crystals required strict maintenance of the temperature at 25 °C. For cryoprotection, the crystals were soaked briefly in the crystallization solution supplemented with 15% *2,3*-butanediol and flash-cooled in the liquid nitrogen. A single-wavelength anomalous dispersion (SAD) data set was collected at the Se absorption peak and processed with *HKL2000* ^43^. Se positions and initial electron density map were calculated using the AutoSol module in *PHENIX* ^44^. The model building and structure refinement were performed by using *COOT ^45^* and *PHENIX*.

